# The CK2 and DBT kinases promote temperature compensation of the *Drosophila* circadian clock via distinct pathways

**DOI:** 10.64898/2026.05.18.725963

**Authors:** Yongliang Xia, Victoria Louis, Patrick Emery

## Abstract

Circadian (∼24 h) rhythms are essential for the survival of most organisms, as they optimize physiology and behavior with the time of day. They are defined by three fundamental properties: they are driven by a self-sustained molecular oscillator, entrained by environmental cues such as light and temperature, and temperature-compensated, whereby circadian period remains close to 24 h over a physiological range of temperatures. The molecular basis of temperature compensation remains incompletely understood. Here, we build on previous studies supporting a conserved and important role for phosphorylation-dependent mechanisms in the control of temperature compensation. We found that reducing the activity of two highly conserved circadian kinases, DBT (casein kinase [CK] 1) and CK2, disrupts temperature compensation in *Drosophila*. Genetic analyses indicate that DBT and CK2 act through distinct pathways that have additive effects on temperature compensation. DBT acts through the *per^Short^* phosphorylation cluster and the S47 phosphodegron of the core clock protein PER, both of which are required for normal thermal compensation. In contrast, CK2 acts through a phosphocluster in TIM as well as PER S45 residue. Interestingly, simultaneous disruption of both pathways causes accumulation of hyperphosphorylated PER, which is inefficiently cleared from the nucleus of circadian pacemaker neurons. Combined with previous work, our findings support a central and unifying role for nuclear PER phosphorylation dynamics in buffering circadian period against environmental temperature fluctuations.

## Introduction

The circadian clock is an endogenous ∼24-hour (h) timekeeping system that enables organisms to anticipate and adapt to predictable daily environmental cycles. By aligning physiological and behavioral processes with the time of day, circadian regulation optimizes energy utilization and maintains internal homeostasis^1–3^. In eukaryotes, the molecular architecture of circadian clocks is centered on interlocked transcriptional feedback loops, with transcriptional activators driving the expression of genes encoding their own repressors^4^, thereby generating self-sustained oscillations. In *Drosophila melanogaster*, for example, the transcription factors CLOCK (CLK) and CYCLE (CYC) form a heterodimer that activates the transcription of *period (per)* and *timeless (tim)*. The PER and TIM proteins accumulate in the cytoplasm, dimerize, and translocate into the nucleus, where they repress CLK–CYC-mediated transcription^5,6^. This negative feedback loop takes approximately 24 hours to complete and constitutes the core oscillator. Proper circadian function requires a balance between plasticity and robustness^7,8^: plasticity enables entrainment to environmental cues such as light–dark cycles, whereas robustness ensures stable timekeeping despite environmental fluctuations (e.g., temperature changes) and internal perturbations (e.g., metabolic variability). Together, these properties allow organisms to maintain precise rhythms in dynamic environments.

Although transcriptional feedback loops form the core of circadian timing, post-translational modifications are equally critical for generating accurate and stable oscillations. Modifications such as phosphorylation, ubiquitination, glycosylation, acetylation and SUMOylation regulate the stability, activity, subcellular localization, and interaction dynamics of clock proteins^9–15^, thereby fine-tuning circadian period and phase. Consistent with this, mutations affecting kinases, phosphatases, or specific modification sites often lead to arrhythmicity or altered free-running periods across species^11,16–18^. Notably, the first identified mammalian circadian mutant, *tau*, carried a mutation in casein kinase Iε (CK1ε)^19^, which alters the degradation kinetics of PERIOD proteins. Similarly, the *Drosophila doubletime* (DBT) kinase, the fly homolog of mammalian CK1ε also regulates PER degradation rates in flies^20–22^.

At its core, the circadian clock operates as a biochemical oscillator driven by cyclic molecular interactions and reaction kinetics. According to the Arrhenius relationship, increasing temperature accelerates biochemical reaction rates and would thus be expected to shorten oscillatory periods. However, circadian clocks exhibit a striking property: their free-running period remains nearly constant across a broad range of physiological temperatures^23,24^. This phenomenon, termed temperature compensation^24^, is one of the defining features of circadian systems, alongside self-sustained oscillation and entrainment^25^. Despite decades of investigation, the molecular basis of temperature compensation remains incompletely understood.

Substantial effort has been devoted to elucidating temperature compensation through both experimental and theoretical approaches. Genetic studies across multiple organisms have identified mutations that disrupt temperature compensation in genes involved in diverse processes, including mRNA 3′-end processing^26^, protein–protein interactions^27^, pre-mRNA metabolism^28^, subcellular protein localization^29,30^, ubiquitin-like modifications^31^, protein methylation^32^, calcium signaling^33^, and the regulation of clock protein stability and activity^34–36^. It remains unclear whether these factors act independently or converge within an integrated regulatory network. More fundamentally, it remains unclear whether a unifying and evolutionarily conserved mechanism underlies temperature compensation across phyla.

Complementing experimental work, a variety of computational models have been proposed to explain temperature compensation. The antagonistic balance model posits that temperature-dependent reactions that accelerate the clock are offset by others that decelerate it^24,37^, resulting in a stable period. Enzyme-limited models instead emphasize competition among reactions for shared, limiting enzymatic resources^38^, providing an intrinsic buffering mechanism. Feedback-based models such as the Goodwin model show that negative feedback best supports temperature compensation and suppresses intrinsic noise, whereas positive feedback reduces extrinsic noise^39^. More detailed transcription–translation models, suggest that temperature compensation emerges from the coordinated behavior of multiple temperature-sensitive processes^40^. Notably, several models converge on the idea that the thermal sensitivity of protein degradation plays a central role in temperature compensation^41–43^.While these models provide valuable conceptual insights, they require further experimental validation and are likely to capture complementary aspects of the underlying biology.

Comparative analyses reveal that core clock genes are not universally conserved, suggesting independent evolution in different kingdoms^4,44^. However, recent studies indicate that circadian rhythms might have originated from an ancient mechanism implicating enzymes with extremely slow NTPase activity^45^. Two fundamental design principles recur across species: rhythmic transcriptional regulation and multisite phosphorylation^4,46,47^. In some organisms, particularly cyanobacteria, temperature compensated circadian rhythms can be driven predominantly by a post-translational oscillator based on rhythmic phosphorylation of the core protein KaiC^48,49^. Phosphorylation is a versatile and reversible modification that modulates protein activity, stability, localization, and interaction networks, enabling rapid and dynamic responses to environmental inputs. In contrast to the divergence of core clock components, protein kinases are highly conserved across taxa. Among these, casein kinase 1 (CK1) and casein kinase 2 (CK2) have emerged as key regulators of circadian timing in organisms ranging from fungi and plants to insects and mammals^22,50–56^. Importantly, both kinases have also been implicated in temperature compensation in several systems^27,35,53,57^. In particular, temperature-dependent substrate and product binding by mammalian CK1δ has been proposed to contribute directly to the molecular mechanism underlying circadian temperature compensation^58^. Given their conserved roles in modulating clock protein phosphorylation and stability, CK1 and CK2 represent strong candidates for mediating shared mechanisms of temperature compensation. Investigating their functions in insects may therefore provide critical insight into whether an evolutionarily conserved molecular basis underlies temperature compensation.

In this study, we investigated the effects of CK2 and DBT, the *Drosophila* ortholog of mammalian CK1δ/ε on temperature compensation in *Drosophila melanogaster*. Our results showed that reducing the activity of the conserved kinases DBT and CK2 disrupts temperature compensation. Genetic analyses indicate that DBT acts through the *per^Short^*phosphocluster and S47 of PER, whereas CK2 acts through PER S45 and a phosphocluster in TIM. Simultaneous disruption of both pathways causes accumulation of hyperphosphorylated PER and impaired nuclear clearance in pacemaker neurons. These findings identify PER phosphorylation dynamics and subcellular localization as key determinants of circadian temperature compensation.

## Materials and Methods

### *Drosophila* stocks and maintenance

The following fly strains were used for this study: *w*^1118^, (*Ck2α^Tik^*, *Ck2α^TikR^*)^54^, *dbt^AR^/+*^59^, (*dbt^L^*, *dbt^S^*)^22^, (*S45A*, *S47A*, *S47D*)^60^, (*tim*^0^*;tim^WT^*, *tim*^0^*;tim^3A^*)^61^, (*tim*^0^*;tim^WT^*, *tim*^0^*;tim^S^*^1404^*^A^*)^62^, *per^S^* ^63^, (*per0;;per^WT^/+*, *per0;per^S^*^151–153^*^A^/*+)^64^. All the flies were reared in plastic vials on standard cornmeal/agar medium at 25°C under 12:12 light/dark cycles unless otherwise specified.

### Behavior recording and Analysis

2-7 old male flies were loaded into glass behavioral tubes and used for examining circadian locomotor activity. Flies were entrained for 3 days under 12:12 LD cycles at 25°C and then released into constant darkness (DD) for at least 7 days at the indicated temperatures. The locomotor activity of single males was monitored with *Drosophila* Activity Monitors system (Trikinetics, Waltham, MA, United States) in Percival I36-LL incubators (Percival Scientific, Perry, IA, United States). Behavioral Data was analyzed by using the FaasX software provided with Dr. F Rouyer lab^65^. 5 days of group activity for each genotype starting from the 2^nd^ day of DD was used to determine the period length. For graphs depicting differences between periods measured at 18 and 29°C, periods were averaged within each independent experimental repeat for each genotype and temperature combinations. Each bar thus represents an average of period average over 3-5 independent experiments.

### Preparation of whole fly head extract

Protein extractions were performed essentially as previously described^66^. Briefly, adult male flies (2-7 days old) were exposed to LD for 3 full days at indicated temperatures. On the 4th day of LD, flies were collected at the indicated time points and frozen on dry ice. The frozen flies were mechanically decapitated and heads were homogenized in ice-cold HE buffer (100 mM KCI/20 mM HEPES/5% glycerol/10 mM EDTA/0.1% Triton X-100/1 mM dithiothreitol/0.5 mM phenylmethylsulfonyl fluoride/1X Halt Protease Inhibitor Cocktail, EDTA-Free (100X) (Thermo Scientific, Cat. No. 87785), pH 7.5), 4 X SDS sample buffer was added and the mixture boiled at 100°C for 6 min. The boiled homogenates were centrifuged with 14000g at 4°C, and supernatant was transferred to new tubes and maintained in -80°C until the analysis for PER and TIM by Western blotting.

### Western Blotting and Antibodies

Equal amounts of WT and mutant supernatant from each time point were resolved by polyacrylamide-SDS gel electrophoresis (PAGE) (5.8% with an acrylamide:bisacrylamide ratio of 29.6:0.4) and transferred to nitrocellulose membrane using Semi-Dry Transfer Cell (Bio-Rad). Membranes were then incubated in 5% nonfat milk (Bio-Rad) for 1 h at room temperature with gentle shaking, incubated with primary antibodies overnight at 4°C with gentle shaking. Blots were then washed with 1X TBST for 1 h, incubated with secondary antibodies for 1 h at room temperature with gentle shaking. Membranes were washed again in 1× TBST before imaging. Primary antibodies used for Western blot were rabbit anti-PER 1:10000, guinea pig anti-TIM 1:10000 and mouse anti-Spectrin 1:5000. The following fluorescent secondary antibodies were used: IRDye 800 Goat anti-rabbit (LI-COR 926-32213, 1:10000), IRDye 680 Donkey anti-guinea pig (LI-COR 926-68077, 1:10000) and IRDye 680 Goat anti-mouse (LI-COR 926-68072, 1:10000). Images were acquired using the Odyssey CLx (LI-COR) and analyzed using the software ImageStudioLite version 5.2.5 (LI-COR) and Fiji (http://fiji.sc). (Densitometry of bands was determined with Fiji (v.2.14.0/1.54f). Data are expressed relative to the loading control (Spectrin) and are from three independent biological replicates.)

### Immunostaining and Image Analysis

Adult male *S45A;;dbt^AR^/+* flies and *w*^1118^ controls were entrained for 3 days under 12:12 LD cycles at 25°C. Ten minutes before the transition to DD, temperature was shifted to either 18°C or 29°C. Brains were then dissected under red light at CT4, CT16, CT20 and CT24. To compare mutant and control flies at equivalent circadian (subjective) phases, the dissection times for *S45A;;dbt^AR^/+* flies were adjusted to account for their longer free-running period, Because phase drift begins after the transition to DD at ZT12, corresponding mutant collection times were calculated using the following equation: (WT time point + 12)/WT period = (mutant time point + 12)/mutant period. For example, at 18°C, wild-type flies exhibited a free-running period close to 24 h, whereas *S45A;;dbt^AR^/+* flies showed a period of approximately 34.5 h. Using this correction, the mutant collection times corresponding to WT CT4, CT16, CT20, and CT24 were calculated as CT11, CT28.3 (DD2, CT4.3), CT34 (DD2, CT10), and CT39.8 (DD2, CT15), respectively. The same approach was used at 29°C. Because the free-running period of *S45A;;dbt^AR^/+* flies at 29°C was approximately 30.2 h, the corresponding mutant sampling times were calculated using the same equation. Whole-mount immunohistochemistry was performed as previously described (Zhang *et al.,* 2010)^67^. Brains were incubated with rabbit anti-PER (1:1500; generous gift from Dr. Ralf Stanewsky) and mouse anti-PDF (1:400; Developmental Studies Hybridoma Bank). Samples were mounted in VECTASHIELD Antifade Mounting Medium (Vector Laboratories, USA) and imaged using a Zeiss LSM 900 confocal microscope with optical sections collected at 1-µm intervals.

Image analysis was performed using ImageJ. For each s-LNv neuron, regions of interest (ROIs) corresponding to the nucleus and cytoplasm were manually defined, and the integrated fluorescence intensity of each ROI was measured. The ratio of nuclear to cytoplasmic PER signal was then calculated for each neuron.

## Results

### CK2 and DBT are essential for temperature compensation in *Drosophila*

To determine whether CK2 and DBT contribute to temperature compensation in *Drosophila*, we measured free-running circadian period lengths in loss-of-function (LoF) mutants across three temperatures (18°C, 25°C, and 29°C) under constant darkness (DD). As expected, wild-type (WT) flies exhibited minimal variation in period length across this temperature range, consistent with effective temperature compensation.

We first examined mutations affecting the catalytic subunit of CK2 (CK2α), using two alleles, *Tik* (Timekeeper) and *TikR*^54^. Both mutations alter residues within or proximal to the ATP-binding pocket and severely impair kinase activity. Because homozygous mutants are lethal, we analyzed heterozygous flies (Fig. 1A). Both *Ck2α^Tik^/+* and *Ck2α^TikR^/+* displayed a clear undercompensation phenotype (Fig. 1B and Table S1), characterized by a progressive shortening of the circadian period with increasing temperature. Specifically, the period decreased by approximately 1.2 hours between 18°C and 29°C. Notably, *Ck2α^TikR^*, a spontaneous revertant that largely rescues the dominant long-period phenotype of *Ck2α^Tik^*, failed to restore normal temperature compensation. This dissociation indicates that temperature compensation is not simply a function of baseline period length.

**Fig. 1.**
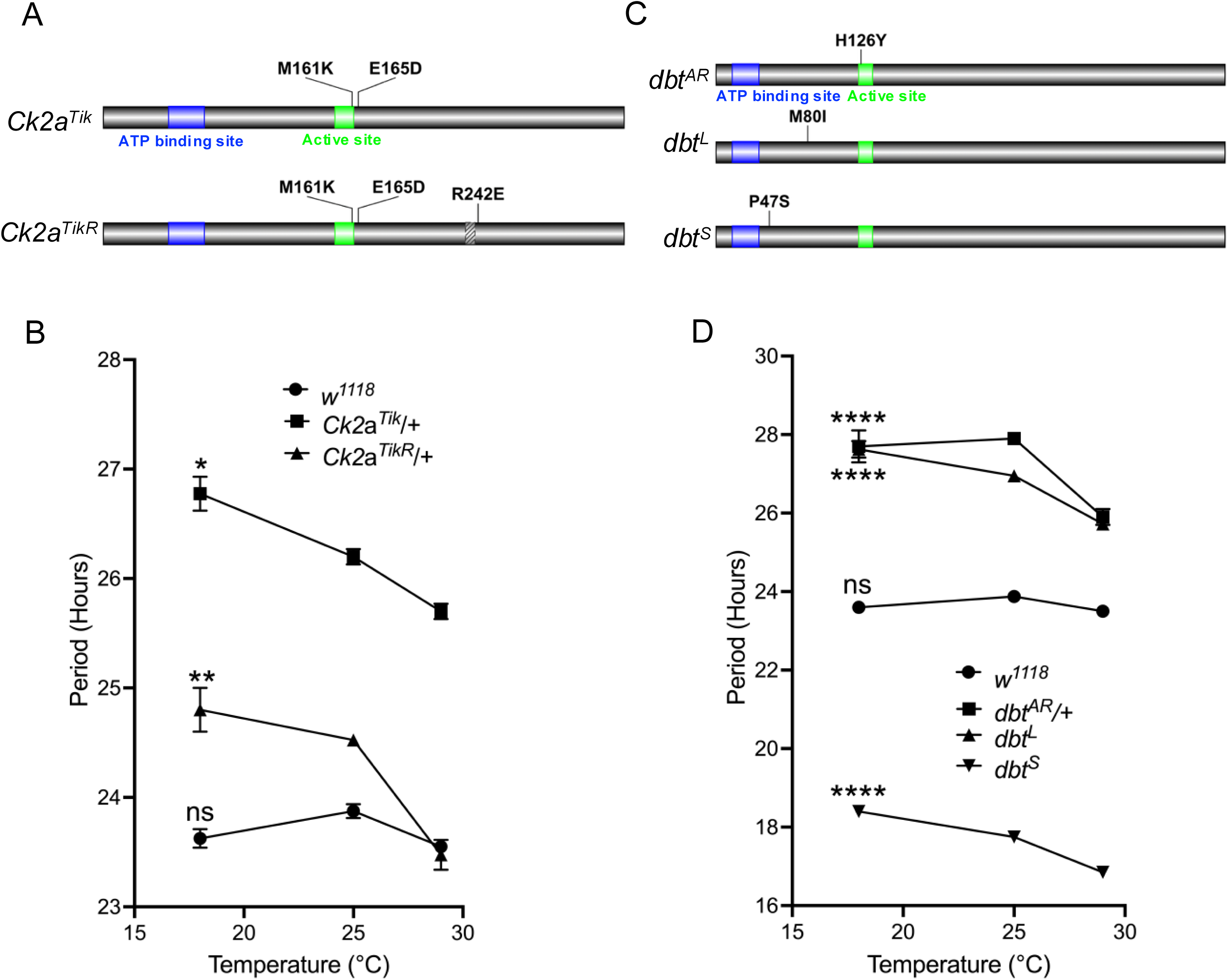
CK2 and DBT influence temperature compensation. **(A)** Schematic of the CK2α protein showing the amino acid alterations in the *Tik* and *TikR* mutants. The slash mark in *TikR* indicates deletion of amino acids 234–240. **(B)** Period lengths of *w*^118^, *Ck2a^Tik^/+* and *Ck2a^TikR^/+* male flies at 18, 25, and 29°C. N = 4 independent experiments. **(C)** Schematic of the DBT protein showing the amino acid alterations in the *dbt^AR^*, *dbt^L^*, and *dbt^S^* mutants. **(D)** Period lengths of *w*^118^, *dbt^AR^/+*, *dbt^L^*and *dbt^S^* male flies at 18, 25, and 29°C. N = 4 independent experiments. Data are presented as mean ± SEM. Statistical significance of differences in free-running period between 18°C and 29°C within each genotype was determined by two-way ANOVA with Sidak’s multiple-comparisons test. *p <0.05; **p < 0.01; ****p < 0.0001; ns, not significant. See also Table S1.

We next assessed the role of DBT, a casein kinase 1δ/ε homolog, using multiple alleles with distinct biochemical properties (Fig. 1C). Heterozygous flies carrying the catalytically impaired allele *dbt^AR^* ^59^ also exhibited undercompensation, with period lengths of approximately 27.8 hours at 18°C and ∼26.0 hours at 29°C (Fig. 1D and Table S1). To further test whether this effect depends on baseline period length, we examined two hypomorphic homozygous alleles, *dbt^S^* (short-period) and *dbt^L^* (long-period)^22^. Despite their opposing effects on period at standard temperature, both mutants consistently showed undercompensation across the tested temperature range, with period shortening of ∼1.5–2 hours from 18°C to 29°C (Fig. 1D and Table S1). Note that both mutants show reduced kinase activity^68^.

Collectively, these results demonstrate that both CK2 and DBT kinase activities are required for robust temperature compensation in *Drosophila*. Importantly, the consistent undercompensation observed across alleles with divergent baseline periods indicates that temperature compensation is mechanistically separable from the determination of absolute circadian period length.

### PER phosphorylation at S47 and the *per^Short^* phosphocluster mediate DBT-dependent temperature compensation

We next sought to identify downstream substrates through which CK2 and DBT regulate temperature compensation. Both kinases are known to phosphorylate the core clock protein PER, thereby modulating its stability, subcellular localization, and transcriptional repressor activity^69–71^. In particular, PER phosphorylation influences its degradation kinetics and nuclear entry^65,72–76^, two processes that are critical for circadian timing.

Previous studies have identified two phosphorylation clusters within PER that are implicated in temperature compensation: the N-terminal phosphodegron (residues S44–S47) and the so-called *per^Shor^*^t^ region (amino acids 583–596)^60^. Both regions contain sites targeted directly by DBT^74^. Within the *per^Shor^*^t^ cluster, three residues (T583, S585, and S589) are phosphorylated by DBT, while S596 is phosphorylated by the kinase NEMO (NMO). Phosphorylation at S596 primes subsequent DBT-dependent phosphorylation at the neighboring residues^74^, demonstrating a hierarchical mechanism. Mutations at three of the four phosphorylation sites result in shortened circadian periods and undercompensation^60,74^, indicating that this cluster functions as a coordinated regulatory module.

To test whether DBT regulates temperature compensation through the *per^Short^* cluster, we analyzed circadian behavior in *per^S^;;dbt^AR^/+* double mutants (Fig. 2A-2A’ and Table S1). The classic *per^S^* allele carries a missense mutation at S589 (S589N), resulting in a short free-running period (∼19.2 h at 25°C) and an undercompensation phenotype (∼1.3 h change from 18°C to 29°C)^77^. Introducing *dbt^AR^/+* into this background lengthened the period by approximately 3 hours at 25°C relative to *per^S^* alone. However, the magnitude of temperature-dependent period change remained essentially unchanged (∼1.3 h), indicating no additive effect of the two mutations on temperature compensation. These findings suggest that DBT controls temperature compensation through the S589 site, and more broadly the *per^Short^* phosphorylation cluster.

**Fig. 2.**
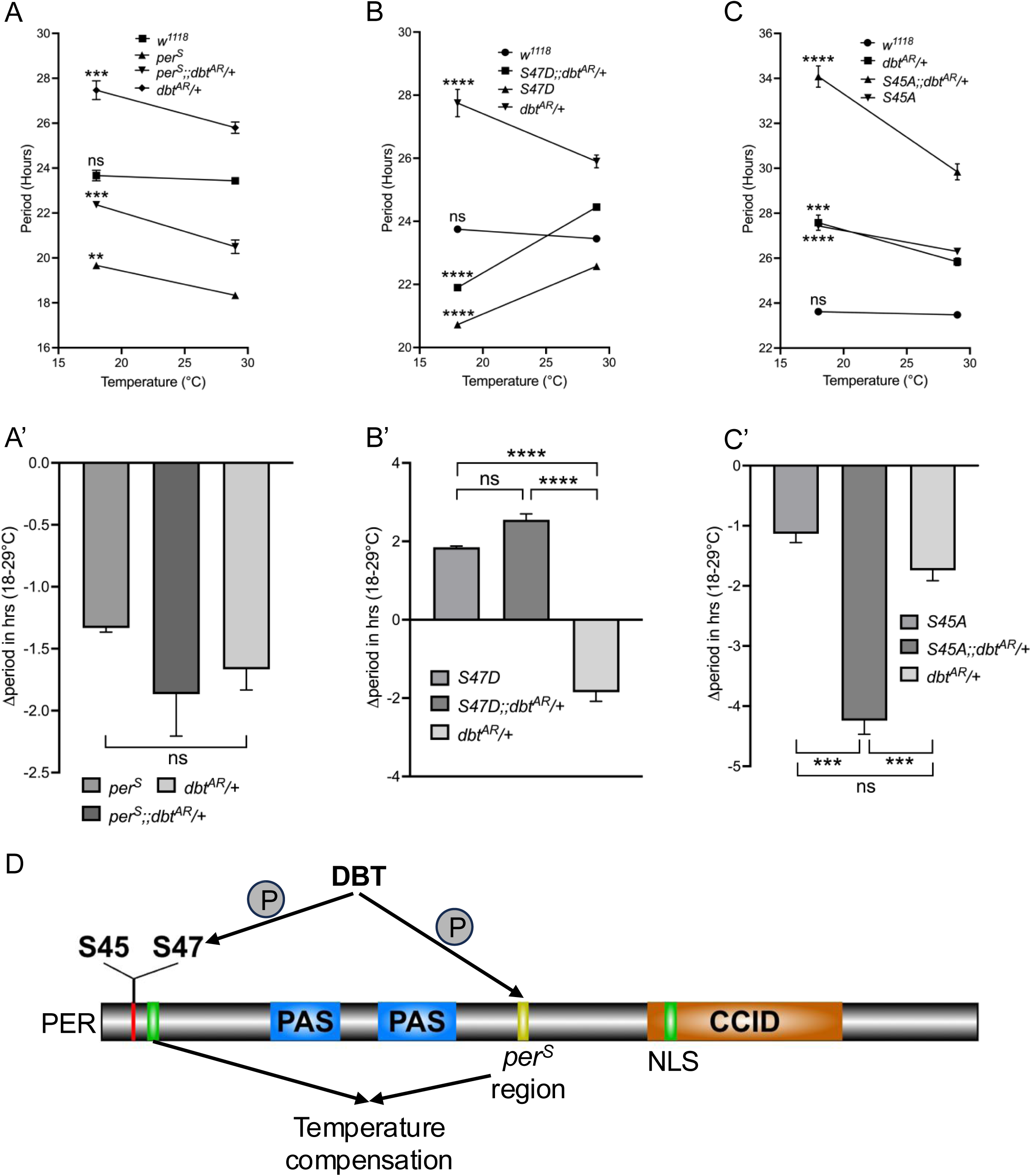
DBT regulates temperature compensation through PER’s S47 phopshodegron and the *per^S^* phpospocluster. **(A)** Period lengths of *w*^118^, *per^S^*, *per^S^;;dbt^AR^/+* and *dbt^AR^/+* male flies at 18 and 29°C. **(A**′) Comparison of the change in period between 18°C and 29°C for each mutant shown in (A). N = 3 independent experiments. **(B)** Period lengths of *w*^118^, *S47D*, *S47D;;dbt^AR^/+* and *dbt^AR^/+* male flies at 18 and 29°C. **(B**′) Comparison of the change in period between 18°C and 29°C for each mutant shown in (B). N = 4 independent experiments. **(C)** Period lengths of *w*^118^, *S45A*, *S45A;;dbt^AR^/+* and *dbt^AR^/+* male flies at 18 and 29°C. **(C**′) Comparison of the change in period between 18°C and 29°C for each mutant shown in (C). N = 5 independent experiments. **(D)** Schematic summary of DBT phosphorylation sites on PER involved in temperature compensation. For (A–C), statistical significance was determined by two-way ANOVA with Sidak’s multiple-comparisons test. For (A′–C′), one-way ANOVA with Tukey’s post hoc test was used. Data are presented as mean ± SEM. **, P<0.01; ***, P<0.001; ****, P<0.0001; ns, not significant. See also Table S1 and Figure S1.

We then examined the contribution of the PER phosphodegron, whose phosphorylation is inhibited by the *per^Short^* domain^74^. Among the three candidate residues in this region, S47 has been experimentally validated as a direct DBT phosphorylation site, and the terminal site in the phosphorylation cascade initiated at S596 by NMO. Phosphorylation at S47 regulates PER stability by modulating its interaction with the F-box protein SLIMB^76^. Consistent with this role, a phosphonull mutation (S47A) lengthens the circadian period, whereas a phosphomimetic mutation (S47D) shortens it^60,76^. Despite their opposite effects on baseline period, both mutants exhibited similar temperature compensation defects, with period increases of approximately 2 hours between 18°C and 29°C^60^.

To determine whether DBT acts through S47 to control temperature compensation, we generated double mutants combining S47 variants with *dbt^AR^/+*. Flies carrying the *S47A;;dbt^AR^/+* genotype were arrhythmic at both 18°C and 29°C, precluding further analysis. In contrast, *S47D;;dbt^AR^/+* flies remained rhythmic and displayed an overcompensation phenotype of ∼2 hours (Fig. 2B-2B’ and Table S1), comparable to that observed in *S47D* single mutants. This epistatic relationship shows that S47D overrides the effect of reduced DBT activity on temperature compensation. Given that S47 is a direct DBT target, these results support the conclusion that DBT regulates temperature compensation predominantly through phosphorylation of S47.

We further examined whether an adjacent residue within the phosphodegron, S45, contribute to DBT-mediated temperature compensation. These residues are thought to cooperate in regulating PER–SLIMB interactions. Mutations at either site result in long-period phenotypes and undercompensation^60^. However, *S45A;;dbt^AR^/+* double mutants exhibited additive effects on both period length and temperature compensation defects (Fig. 2C-2C’ and Table S1), arguing against a shared pathway. This suggests that S45 does not play a significant role in DBT-dependent regulation of temperature compensation.

Taken together, these results indicate that DBT modulates temperature compensation through multiple phosphorylation sites on PER, including S47 within the phosphodegron and at least S589 within the *per^Short^* cluster (Fig. 2D). These findings highlight a central role for PER phosphorylation dynamics in linking kinase activity to temperature-compensated circadian timing.

### PER S45 residue and TIM ST-region phosphorylation mediate CK2-dependent temperature compensation

Given that the S45 residue within the PER phosphodegron contributes to temperature compensation but does not appear to mediate DBT-dependent effects, we next asked whether S45 instead functions downstream of CK2. To test this possibility, we measured circadian period lengths in *S45A;;Ck2α^Tik^/+* and *S45A;;Ck2α^TikR^/+* double mutants at 18°C and 29°C (Fig. 3A, 3A’ and Table S1). Both genotypes exhibited additive effects on baseline period length, particularly in *S45A;;Ck2α^Tik^/+* flies. In contrast, the magnitude of temperature-dependent period change was not further enhanced relative to the single mutants (Fig. 3A’). The absence of additivity in temperature compensation defects indicates that S45 and CK2 act within the same pathway, consistent with a role for S45 in CK2-mediated temperature compensation.

**Fig. 3.**
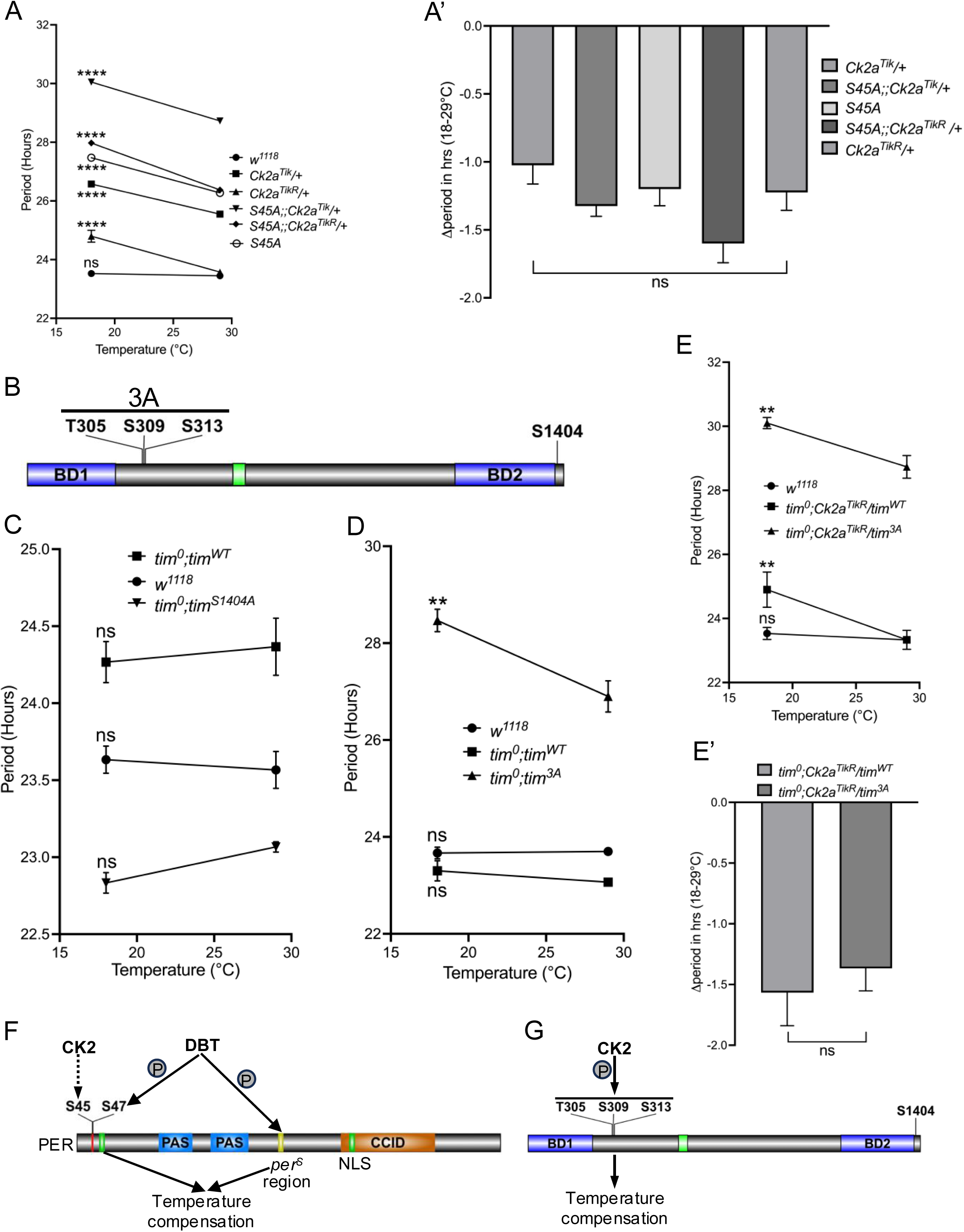
PER S45 and the TIM ST region contribute to CK2-mediated temperature compensation. **(A)** Period lengths of *w*^1118^, *S45A*, *Ck2a^Tik^/+*, *Ck2a^TikR^/+*, *S45A;;Ck2a^Tik^/+* and *S45A;;Ck2a^TikR^/+* male flies at 18 and 29°C. **(A’)** Comparison of the change in period between 18°C and 29°C for each mutant shown in (A). N = 4 independent experiments. **(B)** Schematic of the TIM protein showing CK2 phosphorylation sites. **(C)** Period lengths of *w*^1118^, *tim*^0^*;tim^WT^*, *tim*^0^*;tim^S^*^1404^*^A^*male flies at 18 and 29°C. N = 3 independent experiments. **(D)** Period lengths of *w*^1118^, *tim*^0^*;tim^WT^*, *tim*^0^*;tim^3A^* male flies at 18 and 29°C. N = 3 independent experiments. **(E)** Period lengths of *w*^1118^, *tim*^0^*; Ck2a^TikR^/tim^WT^* and *tim*^0^*;Ck2a^TikR^/tim^3A^*male flies at 18 and 29°C. **(E’)** Comparison of the change in period between 18°C and 29°C for each mutant shown in (E). N = 3 independent experiments. **(F–G)** Schematic summary of phosphorylation sites on PER (F) and TIM (G) involved in CK2-mediated temperature compensation. The dotted arrow indicates that CK2-mediated temperature compensation through PER S45 may occur indirectly. For (A) and (C–E), statistical significance was determined by two-way ANOVA with Sidak’s multiple-comparisons test. For (A’), one-way ANOVA with Tukey’s post hoc test was used. For (E’), an unpaired two-tailed Student’s *t*-test was used. Error bars represent SEM. **, P<0.01; ****, P<0.0001; ns, not significant. See also Table S1.

We next examined additional PER residues reported to be phosphorylated by CK2, specifically S151 and S153. Mutations at these sites delay PER nuclear entry and lengthen circadian period, phenocopying reduced CK2 activity^64^. However, *per*^0^ flies rescued with a *per^S^*^151–153A^ transgene displayed normal temperature compensation across the tested temperature range (Fig. S1), comparable to genetic controls. These observations show that, despite their role in period determination, S151 and S153 are unlikely to mediate CK2-dependent temperature compensation.

In addition to PER, both CLK and TIM have been identified as CK2 substrates. CK2-dependent phosphorylation of CLK has been reported to increase protein stability while repressing its transcriptional activity^78^. However, because the relevant phosphorylation sites on CLK have not been definitively mapped, their contribution to temperature compensation could not be directly assessed. We therefore focused on TIM, for which defined CK2 phosphorylation sites are available (Fig. 3B). One such site, S1404, regulates TIM nuclear export: phosphorylation at this residue inhibits interaction with the nuclear export factor dCRM1, whereas a non-phosphorylatable substitution (S1404A) shortens the circadian period^62^. Despite this effect on baseline period*, tim*^0^*;tim^S^*^1404^*^A^* flies maintained stable period lengths between 18°C and 29°C (Fig. 3C and Table S1), indicating intact temperature compensation and arguing against a role for this site in CK2-dependent temperature compensation.

We next examined a conserved serine/threonine-rich (ST) region of TIM containing three CK2-targeted residues (T305, S309, and S313). Together with SGG-dependent phosphorylation, CK2-mediated modification of this region regulates nuclear accumulation of the PER/TIM complex in small ventral lateral neurons (s-LNvs)^61^. Flies carrying phospho-null substitutions at these sites (3A: T305A/S309A/S313A) exhibited as previously reported a long-period phenotype^61^, consistent with delayed nuclear entry of the PER/TIM complex. Notably, these mutants also displayed pronounced undercompensation, with period lengths of ∼28 hours at 18°C and ∼26 hours at 29°C (Fig. 3D and Table S1), indicating a substantial temperature-dependent shortening of the circadian cycle. Moreover, no additive effects on temperature compensation were observed when combining the 3A mutations with *Ck2α^TikR^* (Fig. 3E, 3E’ and Table S1).

Collectively, these results strongly suggest that CK2 contributes to temperature compensation through multiple amino acid residues, including S45 within the PER phosphodegron and the ST phosphorylation cluster of TIM (Fig. 3F-G). At the present time, there is no evidence that CK2 phosphorylates S45. Our previous work actually suggested a structural role for this amino acid residue, rather than being a phosphorylation target^60^ (see also discussion).

### Simultaneous disruption of the DBT and CK2 pathways sensitize PER phosphorylation to changes in temperature

To further investigate the mechanisms by which DBT and CK2 regulate temperature compensation, we analyzed molecular clock dynamics in *S45A;;dbt^AR^/+* double mutants at 18°C and 29°C. Among the genotypes examined, these flies exhibit the most pronounced undercompensation phenotype, with the circadian period shortening by approximately 4.3 hours across this temperature range (Fig. 2C and Table S1).

We first assessed the temporal profiles of PER and TIM abundance by sampling at six Zeitgeber time (ZT) points under 12:12 light–dark (LD) cycles. Compared with wild-type controls, PER protein levels were consistently elevated in *S45A;;dbt^AR^/+* mutants at both temperatures (Fig. 4A-B and 4D-E), with the most pronounced differences observed during the light phase. In contrast, the overall TIM oscillatory pattern was largely preserved at 18°C (Fig. 4A and 4C), although it appeared to be delayed and reduced amplitude. These phenotypes would be expected given the long period phenotype of these double-mutant animals, and PER’s poor overall abundance rhythms. Moreover, quantitative differences were evident at 29°C: TIM levels were significantly increased at ZT4 and reduced at ZT16 relative to WT (Fig. 4D and 4F), resulting in a diminished oscillation amplitude. Again, the phase of TIM cycle was delayed.

**Fig. 4.**
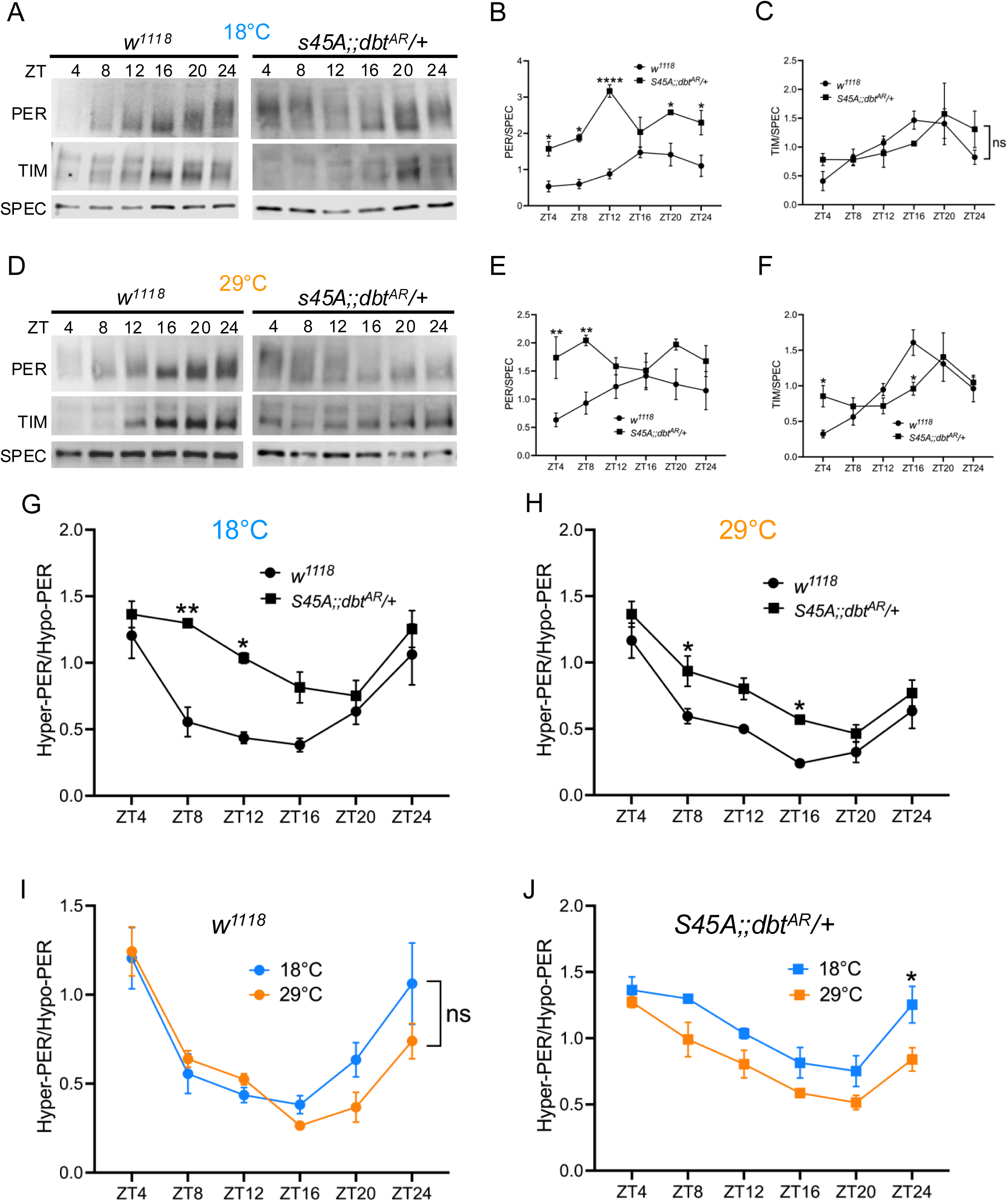
PER abundance and phosphorylation rhythms are disrupted in *S45A;;dbtAR/+* mutants. **(A)** Representative western blots showing temporal PER and TIM expression profiles in head extracts from *w*^118^ and *S45A;;dbt^AR^/+* flies entrained under LD cycles at 18°C. SPEC, spectrin. **(B–C)** Quantification of temporal PER (B) and TIM (C) abundance from flies entrained under LD cycles at 18°C. **(D)** Representative western blots showing temporal PER and TIM expression profiles in head extracts from *w*^118^and *S45A;;dbt^AR^/+* flies entrained under LD cycles at 29°C. **(E–F)** Quantification of temporal PER (E) and TIM (F) abundance from flies entrained under LD cycles at 29°C. **(G–H)** Ratio of hyperphosphorylated PER to hypophosphorylated PER in *w*^118^and *S45A;;dbt^AR^/+* flies at 18°C (A) and 29°C (B). **(I–J)** Ratio of hyperphosphorylated PER to hypophosphorylated PER at 18°C versus 29°C in *w*^118^ (C) and *S45A;;dbt^AR^/+* (D) flies. Error bars represent SEM. Statistical significance was determined by two-way ANOVA with Tukey’s post hoc test. *p <0.05; **p < 0.01; ****p < 0.0001; ns, not significant.

In addition to abundance rhythms, PER in WT flies exhibits robust daily changes in phosphorylation state^79^. To determine whether these dynamics were altered in the mutants, we quantified the ratio of hyperphosphorylated to hypophosphorylated PER at both temperatures. Consistent with the increase in total PER levels, *S45A;;dbt^AR^/+* mutants displayed elevated levels of hyperphosphorylated PER at both 18°C and 29°C (Fig. 4G-H), particularly between ZT8 and ZT16. This persistence of hyperphosphorylated PER explains the poor rhythmicity of total PER abundance observed in the double-mutant on Fig 4A-B and 4D-E.

In WT flies, PER phosphorylation rhythms are themselves temperature-compensated (Fig. 4I), showing similar profiles across the 18–29°C range. In contrast, although the temporal pattern of PER phosphorylation cycling in *S45A;;dbt^AR^/+* mutants remained broadly intact, the absolute levels of hyperphosphorylated PER were consistently higher at 18°C than at 29°C across all time points (Fig. 4J), indicating that PER phosphorylation itself is undercompensated. ANOVA analysis indeed confirmed an overall statistically significant effect of temperature on the hyper/hypophosphorylated PER ratio in the double mutant, that was absent in wild-type (Fig. 4I).

Taken together, these results demonstrate that combined disruption of PER S45 (and hence CK2 signaling) and reduced DBT activity leads to the accumulation of hyperphosphorylated PER, elevating overall PER abundance and altering the amplitude of clock protein oscillations. This accumulation is more pronounced at higher temperature. Moreover, the decrease in temperature compensation of PER phosphorylation dynamics parallels the behavioral phenotype, suggesting that dysregulated PER phosphorylation is a key molecular determinant of the undercompensated circadian rhythms observed in *S45A;;dbt^AR^/+* flies.

### Disruption of the DBT and CK2 pathways impacts PER nuclear localization

Both DBT- and CK2-dependent phosphorylation are known to regulate the nuclear accumulation of PER or PER-TIM complex by modulating either nuclear import or nuclear export^80–84^. In addition, a recent study showed that PER nuclear export plays an important role in circadian temperature compensation^29^. In particular, the *per^I^*^530^*^A^* mutation, which causes overcompensation, leads to prolonged nuclear retention of PER during the day at elevated temperature. We therefore asked whether the *S45A;;dbt^AR^/+* double-mutation impacts PER subcellular localization in the small ventral Lateral neurons (s-LNvs), the pacemaker neurons that drive circadian locomotor behavior in DD^85,86^. To address this question, we stained brains from flies entrained under LD cycles and then released into DD under either 18°C or 29°C. Flies were collected during the first subjective day and the following subjective night at four circadian time points: CT4, CT16, CT20 and CT24 (Fig. 5A). These time points capture key stages of PER re-localization between the cytoplasm and nucleus^87^. To allow comparison of mutant and control flies at equivalent circadian phases, the timing of dissections for *S45A;;dbt^AR^/+* flies was adjusted to account for their long-period phenotype (see Methods for details).

**Fig. 5.**
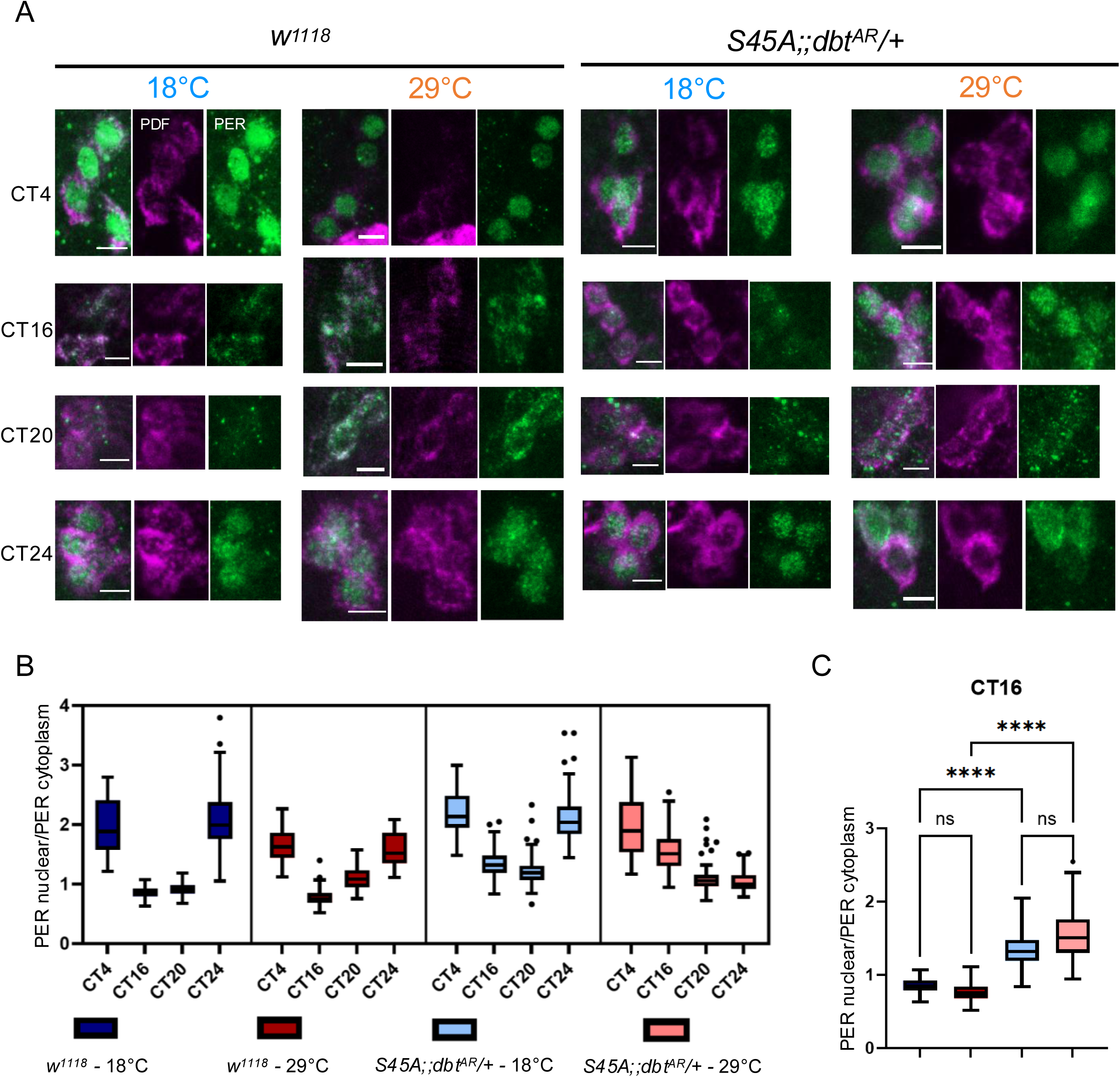
*S45A;;dbt^AR^/+* alters PER subcellular localization in s-LNvs in a temperature-dependent manner. (A) Representative images of PER and PDF staining in s-LNv neurons in control (*w*^118^) and *S45A;;dbt^AR^/+* flies at 18°C and 29°C. Images show maximum-intensity projections of Z-stacks. Scale bar, 5 μm. (B) Ratio of nuclear to cytoplasmic PER signal across circadian time in control and *S45A;;dbt^AR^/+* flies at 18°C and 29°C. (C) Comparison of the nuclear to cytoplasmic PER ratio at CT16 (Kruskal-Wallis test; ****p<0.0001; ns, not significant). See also Figure S2.

Two independent experiments were performed, which yielded similar patterns of rhythmic PER subcellular localization (Fig.5 and S2). In wild-type controls, as expected, PER staining was predominantly nuclear at CT4 and CT24, whereas at CT16 and CT20 PER was mainly cytoplasmic or evenly distributed between the cytoplasm and nucleus (Fig. 5A-B and S2. A). This pattern was observed at both 18°C and 29°C, indicating that the progression of PER subcellular localization during the circadian cycle is itself temperature compensated. In *S45A;;dbt^AR^/+* flies, however, we observed excessive PER nuclear accumulation at CT16 at both temperatures (Fig. 5B-C and S2. A-B), and the rhythm of PER subcellular localization was delayed at warm temperature. Together, these results indicate that nuclear PER is inefficiently cleared from the nucleus in the double mutant, reminiscent of the observations made with *per^I^*^530^*^A^* mutants^29^.

## Discussion

A central property of circadian clocks is temperature compensation, the ability to maintain a nearly constant period over a range of physiological temperatures despite the intrinsic temperature dependence of most biochemical reactions. Although this phenomenon has been recognized for decades^23^, the molecular mechanisms that stabilize clock pace across temperature remain incompletely understood. Our results identify the kinases CK2 and DBT as key components of this regulatory system in *Drosophila* melanogaster. Reduced activity of either kinase caused robust undercompensation (Fig. 1B and 1D), with circadian period shortening as temperature increased. Importantly, this defect was observed across alleles with opposite effects on baseline period length, indicating that temperature compensation is not simply a secondary consequence of altered clock speed but reflects a genetically separable mechanism. This conclusion is consistent with earlier studies showing that mutations affecting temperature compensation can be dissociated from those that determine absolute circadian period^88–90^. Consistent with this idea, a recent study in *Drosophila melanogaster* showed that mutations affecting the interaction interface between PER and DBT consistently lengthened circadian period, yet produced distinct temperature compensation phenotypes^91^.

Our genetic analyses further indicate that temperature compensation emerges from coordinated phosphorylation of multiple clock substrates rather than from regulation of a single rate-limiting step. For PER, we identified two DBT-dependent modules that contribute to thermal robustness: S47 within the phosphodegron and S589 within the *per^Short^* cluster (Fig. 2). Both regions have previously been implicated in PER stability and degradation kinetics^74,76^. The absence of additive temperature compensation defects in the relevant double mutants suggests that these residues function within a common DBT-dependent pathway. By contrast, S45 in the PER phosphodegron behaved differently. Although mutation of S45 altered temperature compensation, its interaction with *dbt* was additive (Fig. 2C and 2C’), whereas no additivity was observed with reduced CK2 activity (Fig. 3A and 3A’). These results place S45 functionally within the CK2 pathway. Notably, other CK2-dependent PER residues previously shown to regulate nuclear entry, including S151 and S153, did not affect temperature compensation (Fig. S1), indicating that only a subset of CK2-dependent phosphorylation events participates in thermal buffering.

Our data further identify TIM as an additional relevant target of CK2. Phosphorylation of the TIM serine/threonine-rich region, previously implicated in regulating nuclear accumulation of the PER/TIM complex in clock neurons^61^, proved necessary for normal temperature compensation (Fig. 3D), whereas mutation of TIM S1404 had no detectable effect (Fig. 3C). These observations support a model in which CK2 acts through multiple substrates (Fig. 3F-G), rather than exclusively through PER. Such distributed regulation may provide robustness by allowing temperature-dependent changes in one molecular process to be offset by compensatory effects at another.

The molecular analyses of the strongly undercompensated *S45A;;dbt^AR^/+* double mutant provide insight into how these phosphorylation pathways stabilize circadian timing. Combined disruption of CK2- and DBT-dependent signaling caused elevated PER abundance, persistent accumulation of hyperphosphorylated PER, and reduced amplitude of PER and TIM oscillations (Fig. 4A-H). Importantly, whereas wild-type flies maintained nearly identical PER phosphorylation profiles at 18°C and 29°C, the double mutant exhibited a significant temperature-dependent reduction in the hyper-phosphorylated/hypo-phosphorylated PER ratio (Fig. 4I-J), which echoes its behavioral undercompensation phenotype. Thus, the biochemical temperature compensation of PER phosphorylation itself is compromised in the mutant. These findings argue that one critical function of DBT and CK2 is to maintain the balance among PER phospho-isoforms across temperature. This interpretation is consistent with prior work linking PER phosphorylation dynamics to temperature-dependent control of circadian period^60,92–94^.

Our results further support the idea that, in addition to its phosphorylation^60^, the regulation of PER subcellular localization is critical for temperature compensation^29^. Interestingly, Giesecke et al. observed that the overcompensated *per^I^*^530^*^A^* mutant retains PER in the nucleus for prolonged periods at warm temperatures. We observed a related phenotype in *S45A;;dbt^AR^/+* flies, where PER also accumulated abnormally in the nucleus and was inefficiently cleared during subjective day (Fig. 5 and S2). At first glance, the fact that both mutants exhibit excessive nuclear PER despite opposite behavioral phenotypes, overcompensation in *per^I^*^530^*^A^* and undercompensation in *S45A;;dbt^AR^/+* appears paradoxical. However, there is a crucial biochemical distinction between the two conditions. In *per^I^*^530^*^A^*, PER remains predominantly hypo-phosphorylated, whereas in *S45A;;dbt^AR^/+* we observed pronounced accumulation of hyperphosphorylated PER (Fig. 4G-H). We therefore propose that nuclear hypophosphorylated PER promotes temperature compensation, whereas nuclear hyperphosphorylated PER antagonizes it. In this model, the relevant determinant is not simply nuclear residence time, but the phosphorylation state of nuclear PER. DBT and CK2, together with PER degradation pathways, would therefore maintain the ratio of hyper- and hypo-phosphorylated nuclear PER within a narrow range required for precise temperature compensation (Fig. 6).

**Fig. 6.**
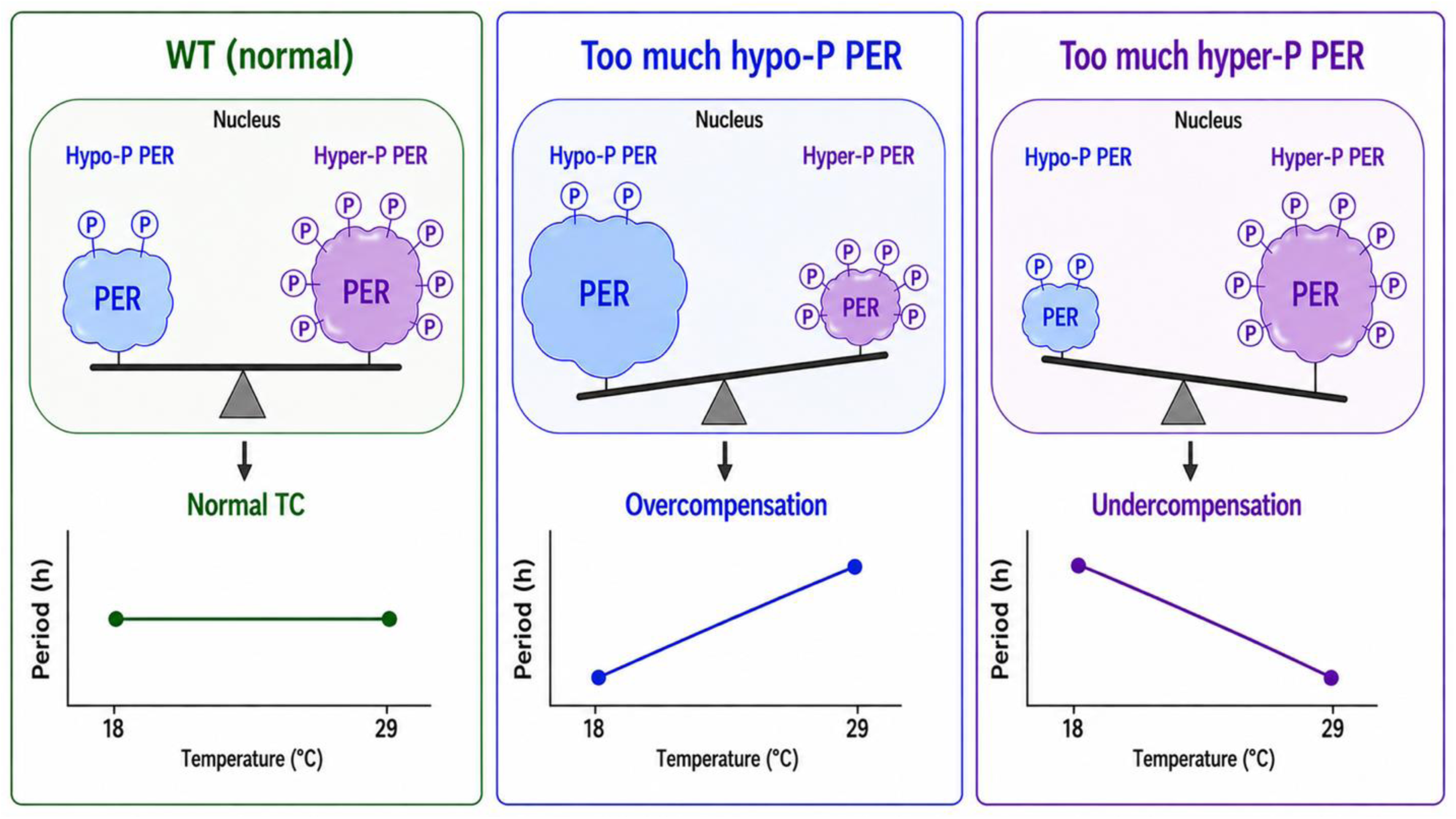
Working model: the phosphorylation state of nuclear PER determines circadian temperature compensation. The critical determinant of temperature compensation is not simply the duration of nuclear PER residence, but the phosphorylation state of nuclear PER. In wild-type flies, coordinated activities of DBT (CKI), CK2, and PER degradation pathways maintain the balance between hypo- and hyperphosphorylated nuclear PER within a narrow range, thereby enabling precise temperature compensation. Mutations that shift this balance toward hypophosphorylated nuclear PER promote overcompensation, whereas mutations that favor hyperphosphorylated nuclear PER cause undercompensation.

This model provides a framework that reconciles apparently disparate observations. For example, mutations that affect distinct phosphorylation sites or reduce kinase activity produce very different baseline circadian periods, yet lead to similar temperature-compensation defects. This is the case for the different CK2α and DBT alleles examined in this study. These findings argue that temperature compensation is not determined by the phosphorylation state of any single residue, by PER stability alone, or by baseline circadian period length. Instead, our results suggest that precise temperature compensation depends on maintaining the proper balance and distribution of PER phospho-states within the circadian system, particularly in the nucleus. Perturbations that shift this balance toward either excessively hypophosphorylated or hyperphosphorylated nuclear PER impair thermal compensation, despite producing distinct molecular and behavioral phenotypes. Temperature compensation may therefore emerge from dynamic control of PER phospho-state distribution rather than from simple modulation of mean kinase activity.

Several limitations of our study should be noted. First, it is not yet clear whether S45 is a direct substrate of CK2 or whether it acts indirectly through structural effects on PER or PER/TIM complex conformation. Our previous study suggests that S45 more likely functions structurally rather than being modulated by phosphorylation, at least as far as temperature compensation is concerned^60^. Second, CK2 might act on additional substrates, including CLK, whose contribution could not be addressed here because the relevant phosphosites remain incompletely defined. Third, our biochemical analyses were performed in head extracts and therefore reflect the summed PER phosphorylation pattern in multiple tissues (predominantly the eyes), rather than only the circadian pacemaker neurons. Cell-type-specific analyses of phosphorylation dynamics and nuclear trafficking would be needed to determine how these mechanisms operate within the pacemaker neurons that control free-running locomotor rhythms.

In summary, our findings support a model in which temperature compensation depends on coordinated control of PER phosphorylation, degradation, and subcellular localization. By acting on multiple sites in PER and TIM, CK2 and DBT buffer the circadian oscillator against temperature-dependent changes in reaction rates and preserve stable circadian timing. More broadly, these results support the model that temperature compensation is an emergent property of balanced phosphoregulatory network dynamics within the core clock.

## Data availability

All data necessary for supporting the findings of this study are included within the article and its accompanying figures and tables. Any additional materials or information described in this manuscript are available from the corresponding authors upon request.

## Acknowledgments

We thank Joanna C. Chiu, Deniz Top, Jan Pielage, and Ravi Allada for generously providing *Drosophila* lines used in this study. We also acknowledge the Bloomington Drosophila Stock Center for supplying additional fly stocks, the center is supported by the NIH Office of Research Infrastructure Programs (P40 OD018537). We thank Lauren North and Vinh Phan for assistance with fly rearing and stock maintenance. Finally, we are grateful to all members of the laboratory for their valuable discussions, constructive feedback, and continued support throughout the course of this project.

## Figure legends

**Fig. S1. Period lengths of *w***^1118^**, *per***^0^***;per^WT^/+*, *per***^0^***;per^S^***^151–153^***^A^/+* male flies at 18 and 29°C.** N = 3 independent experiments. Data are presented as mean ± SEM. Statistical significance of differences in free-running period between 18°C and 29°C within each genotype was determined by two-way ANOVA with Sidak’s multiple-comparisons test. ns, not significant. See also Table S1. Related to Figure 2.

**Fig. S2. *S45A;;dbt^AR^/+* alters PER subcellular localization in s-LNvs in a temperature-dependent manner.** (A) Ratio between nuclear to cytoplasmic PER across circadian time in s-LNv neurons of control (*w*^1118^) and *S45A;;dbt^AR^/+* flies at 18°C and 29°C. (B) Comparison of nuclear to cytoplasmic PER ratio at CT16 (Kruskal-Wallis test; **p<0.01; ***p<0.001; ****p<0.0001). Related to Figure 5.

**Table S1.** Circadian locomotor behavior of control and various mutant flies at 18 and 29°C.

| Genotype | Temperature (°C) | N | Rhythmic flies (%) | Period (h) ± SEM | Power ± SEM |
| --- | --- | --- | --- | --- | --- |
| *w<sup>1118</sup>* | 18 | 155 | 52.9 | 23.7 ± 0.06 | 39.9 ± 2.12 |
|  | 29 | 150 | 94.7 | 23.5 ± 0.02 | 70.6 ± 3.51 |
| *CK2a<sup>Tik</sup>/+* | 18 | 43 | 74.4 | 26.8 ± 0.13 | 58.4 ± 6.99 |
|  | 29 | 41 | 92.7 | 25.6 ± 0.08 | 59.2 ± 1.83 |
| *CK2a<sup>TikR</sup>/+* | 18 | 45 | 86.7 | 24.6 ± 0.03 | 63.9 ± 4.55 |
|  | 29 | 42 | 88.1 | 23.4 ± 0.10 | 73.3 ± 5.56 |
| *dbt<sup>AR</sup>/+* | 18 | 80 | 51.3 | 27.8 ± 0.32 | 37.5 ± 3.24 |
|  | 29 | 78 | 91.0 | 26.0 ± 0.18 | 59.6 ± 1.77 |
| *dbt<sup>S</sup>* | 18 | 27 | 29.6 | 18.4 ± 0.07 | 57.5 ± 12.49 |
|  | 29 | 22 | 40.9 | 16.9 ± 0.03 | 35.1 ± 4.36 |
| *dbt<sup>L</sup>* | 18 | 48 | 25.0 | 27.6 ± 0.21 | 32.4 ± 4.46 |
|  | 29 | 42 | 47.6 | 25.7 ± 0.17 | 53.9 ± 6.65 |
| *per<sup>S</sup>* | 18 | 94 | 77.7 | 19.7 ± 0.02 | 69.1 ± 3.42 |
|  | 29 | 77 | 46.8 | 18.5 ± 0.10 | 60.9 ± 5.89 |
| *per<sup>S</sup>;;dbt<sup>AR</sup>/+* | 18 | 34 | 73.5 | 22.4 ± 0.07 | 49.4 ± 6.66 |
|  | 29 | 34 | 94.1 | 20.5 ± 0.30 | 66.8 ± 10.60 |
| *S47D* | 18 | 86 | 73.3 | 20.8 ± 0.06 | 45.4 ± 2.79 |
|  | 29 | 86 | 80.2 | 22.5 ± 0.07 | 57.1 ± 1.72 |
| *S47D;;dbt<sup>AR</sup>/+* | 18 | 64 | 87.5 | 21.9 ± 0.07 | 53.8 ± 4.03 |
|  | 29 | 63 | 90.5 | 24.5 ± 0.12 | 54.9 ± 1.44 |
| *S45A* | 18 | 51 | 70.6 | 27.5 ± 0.08 | 46.8 ± 2.62 |
|  | 29 | 48 | 91.7 | 26.3 ± 0.05 | 60.2 ± 2.75 |
| *S45A;;dbt<sup>AR</sup>/+* | 18 | 58 | 39.7 | 34.5 ± 0.33 | 33.8 ± 3.10 |
|  | 29 | 57 | 89.5 | 30.2 ± 0.15 | 67.0 ± 3.73 |
| *S45A;;CK2a<sup>Tik</sup>/+* | 18 | 67 | 97.0 | 30.1 ± 0.04 | 67.2 ± 4.26 |
|  | 29 | 64 | 96.9 | 28.7 ± 0.07 | 71.9 ± 3.32 |
| *S45A;;CK2a<sup>TikR</sup>/+* | 18 | 70 | 95.7 | 28.0 ± 0.09 | 75.4 ± 4.07 |
|  | 29 | 66 | 97.0 | 26.4 ± 0.05 | 68.9 ± 2.10 |
| *tim<sup>0</sup>;tim<sup>WT</sup>* | 18 | 52 | 23.1 | 24.2 ± 0.12 | 43.9 ± 4.66 |
|  | 29 | 49 | 18.4 | 24.5 ± 0.16 | 45.6 ± 7.06 |
| *tim<sup>0</sup>;tim<sup>S1404A</sup>* | 18 | 54 | 33.3 | 22.8 ± 0.15 | 53.0 ± 7.25 |
|  | 29 | 50 | 56.0 | 23.2 ± 0.16 | 61.5 ± 11.5 |
| *tim<sup>0</sup>;tim<sup>WT</sup>* | 18 | 60 | 10.0 | 23.4 ± 0.18 | 36.6 ± 11.31 |
|  | 29 | 62 | 91.9 | 23.0 ± 0.05 | 73.9 ± 4.85 |
| *tim<sup>0</sup>;tim<sup>3A</sup>* | 18 | 57 | 38.6 | 28.4 ± 0.19 | 34.0 ± 2.96 |
|  | 29 | 62 | 91.9 | 27.0 ± 0.24 | 64.3 ± 1.36 |
| *tim<sup>0</sup>;CK2a<sup>TikR</sup>/tim<sup>WT</sup>* | 18 | 33 | 63.6 | 24.9 ± 0.55 | 38.0 ± 1.11 |
|  | 29 | 35 | 82.9 | 23.3 ± 0.30 | 57.3 ± 7.17 |
| *tim<sup>0</sup>;CK2a<sup>TikR</sup>/tim<sup>3A</sup>* | 18 | 42 | 61.9 | 30.2 ± 0.23 | 44.9 ± 1.17 |
|  | 29 | 39 | 41.0 | 28.4 ± 0.12 | 42.9 ± 3.06 |
| *per<sup>0</sup>;per<sup>WT</sup>/+* | 18 | 43 | 60.5 | 23.9 ± 0.08 | 39.7 ± 6.45 |
|  | 29 | 48 | 93.8 | 23.9 ± 0.05 | 96.8 ± 6.20 |
| *per<sup>0</sup>;per<sup>S151-153A</sup>/+* | 18 | 69 | 95.7 | 25.0 ± 0.09 | 51.2 ± 3.67 |
|  | 29 | 65 | 87.7 | 24.8 ± 0.13 | 66.3 ± 9.32 |

## Literature cited

1 Martel, J., Ojcius, D. & Young, J. D. Circadian rhythmicity and human health. Biomedical Journal 48 (2025).

2 Ayyar, V. S. & Sukumaran, S. Circadian rhythms: influence on physiology, pharmacology, and therapeutic interventions. Journal of Pharmacokinetics and Pharmacodynamics 48, 321–338 (2021).

3 Dunlap, J. C., Loros, J. J. & DeCoursey, P. J. Chronobiology : biological timekeeping. (Sinauer Associates, 2004).

4 Dunlap, J. C. Molecular bases for circadian clocks. Cell 96, 271–290, doi:10.1016/s0092-8674(00)80566-8 (1999).

5 Hardin, P. E. The circadian timekeeping system of Drosophila. Current biology : CB 15 **17**, R714–722 (2005).

6 Zheng, X. & Sehgal, A. Speed control: cogs and gears that drive the circadian clock. Trends in neurosciences 35 **9**, 574–585 (2012).

7 Hatakeyama, T. S. & Kaneko, K. Reciprocity Between Robustness of Period and Plasticity of Phase in Biological Clocks. Physical review letters 115 **21**, 218101 (2015).

8 Thommen, Q., Pfeuty, B., Corellou, F., Bouget, F.-Y. & Lefranc, M. Robust and flexible response of the Ostreococcus tauri circadian clock to light/dark cycles of varying photoperiod. The FEBS Journal 279 (2011).

9 Honma, S. The mammalian circadian system: a hierarchical multi-oscillator structure for generating circadian rhythm. The Journal of Physiological Sciences 68, 207–219 (2018).

10 Yan, J., Kim, Y. J. & Somers, D. E. Post-Translational Mechanisms of Plant Circadian Regulation. Genes 12 (2021).

11 Zhang, H., Zhou, Z. & Guo, J. The Function, Regulation, and Mechanism of Protein Turnover in Circadian Systems in Neurospora and Other Species. International Journal of Molecular Sciences 25 (2024).

12 Sharma, A., et al. Circadian Rhythm Disruption and Alzheimer’s Disease: The Dynamics of a Vicious Cycle. Current Neuropharmacology 19, 248–264 (2020).

13 Kwon, I., Choe, H. K., Son, G. H. & Kim, K. Mammalian Molecular Clocks. Experimental Neurobiology 20, 18–28 (2011).

14 Weber, F., Zorn, D., Rademacher, C. & Hung, H. C. Post-translational timing mechanisms of the Drosophila circadian clock. FEBS Lett 585, 1443–1449, doi:10.1016/j.febslet.2011.04.008 (2011).

15 Tataroglu, O. & Emery, P. The molecular ticks of the Drosophila circadian clock. Curr Opin Insect Sci 7, 51–57, doi:10.1016/j.cois.2015.01.002 (2015).

16 Lin, J.-M., Schroeder, A. M. & Allada, R. In Vivo Circadian Function of Casein Kinase 2 Phosphorylation Sites in Drosophila PERIOD. The Journal of Neuroscience 25, 11175–11183 (2005).

17 Zheng, X. & Sehgal, A. Probing the Relative Importance of Molecular Oscillations in the Circadian Clock. Genetics 178, 1147–1155 (2008).

18 Philpott, J. M., et al. Casein kinase 1 dynamics underlie substrate selectivity and the PER2 circadian phosphoswitch. eLife 9 (2020).

19 Lowrey, P. L., et al. Positional syntenic cloning and functional characterization of the mammalian circadian mutation tau. Science 288 5465, 483–492 (2000).

20 Syed, S., Saez, L. & Young, M. W. Kinetics of Doubletime Kinase-dependent Degradation of the Drosophila Period Protein*. The Journal of Biological Chemistry 286, 27654–27662 (2011).

21 Kloss, B., et al. The Drosophila clock gene double-time encodes a protein closely related to human casein kinase Iepsilon. Cell 94, 97–107, doi:10.1016/s0092-8674(00)81225-8 (1998).

22 Price, J. L., et al. double-time is a novel Drosophila clock gene that regulates PERIOD protein accumulation. Cell 94, 83–95, doi:10.1016/s0092-8674(00)81224-6 (1998).

23 Pittendrigh, C. S. On temperature independence in the clock system controlling emergence time in Drosophila. Proc Natl Acad Sci U S A 40, 1018–1029, doi:10.1073/pnas.40.10.1018 (1954).

24 Hastings, J. W. & Sweeney, B. M. On the mechanism of temperature independence in a biological clock. Proc Natl Acad Sci U S A 43, 804–811, doi:10.1073/pnas.43.9.804 (1957).

25 Pittendrigh, C. S. Circadian rhythms and the circadian organization of living systems. Cold Spring Harb Symp Quant Biol 25, 159–184, doi:10.1101/sqb.1960.025.01.015 (1960).

26 Schmal, C., et al. Alternative polyadenylation factor CPSF6 regulates temperature compensation of the mammalian circadian clock. PLoS Biol 21, e3002164, doi:10.1371/journal.pbio.3002164 (2023).

27 Hu, Y., et al. FRQ-CK1 Interaction Underlies Temperature Compensation of the Neurospora Circadian Clock. mBio 12, e0142521, doi:10.1128/mBio.01425-21 (2021).

28 Marshall, C. M., Tartaglio, V., Duarte, M. & Harmon, F. G. The Arabidopsis sickle Mutant Exhibits Altered Circadian Clock Responses to Cool Temperatures and Temperature-Dependent Alternative Splicing. Plant Cell 28, 2560–2575, doi:10.1105/tpc.16.00223 (2016).

29 Giesecke, A., et al. A novel period mutation implicating nuclear export in temperature compensation of the Drosophila circadian clock. Curr Biol 33, 336–350.e335, doi:10.1016/j.cub.2022.12.011 (2023).

30 Matsumoto, A., Tomioka, K., Chiba, Y. & Tanimura, T. timrit Lengthens circadian period in a temperature-dependent manner through suppression of PERIOD protein cycling and nuclear localization. Mol Cell Biol 19, 4343–4354, doi:10.1128/MCB.19.6.4343 (1999).

31 Hansen, L. L., van den Burg, H. A. & van Ooijen, G. Sumoylation Contributes to Timekeeping and Temperature Compensation of the Plant Circadian Clock. J Biol Rhythms 32, 560–569, doi:10.1177/0748730417737633 (2017).

32 Jones, M. A., Morohashi, K., Grotewold, E. & Harmer, S. L. Arabidopsis JMJD5/JMJ30 Acts Independently of LUX ARRHYTHMO Within the Plant Circadian Clock to Enable Temperature Compensation. Front Plant Sci 10, 57, doi:10.3389/fpls.2019.00057 (2019).

33 Kon, N., et al. Na(+)/Ca(2+) exchanger mediates cold Ca(2+) signaling conserved for temperature-compensated circadian rhythms. Sci Adv 7, doi:10.1126/sciadv.abe8132 (2021).

34 Maeda, A. E., Matsuo, H., Muranaka, T. & Nakamichi, N. Cold-induced degradation of core clock proteins implements temperature compensation in the Arabidopsis circadian clock. Sci Adv 10, eadq0187, doi:10.1126/sciadv.adq0187 (2024).

35 Portolés, S. & Más, P. The functional interplay between protein kinase CK2 and CCA1 transcriptional activity is essential for clock temperature compensation in Arabidopsis. PLoS Genet 6, e1001201, doi:10.1371/journal.pgen.1001201 (2010).

36 Terauchi, K., et al. ATPase activity of KaiC determines the basic timing for circadian clock of cyanobacteria. Proc Natl Acad Sci U S A 104, 16377–16381, doi:10.1073/pnas.0706292104 (2007).

37 Narasimamurthy, R. & Virshup, D. M. Molecular Mechanisms Regulating Temperature Compensation of the Circadian Clock. Frontiers in Neurology 8 (2017).

38 Hatakeyama, T. S. & Kaneko, K. Generic temperature compensation of biological clocks by autonomous regulation of catalyst concentration. Proceedings of the National Academy of Sciences 109, 8109–8114 (2011).

39 Chakravarty, S., Hong, C. I. & Csikász-Nagy, A. Systematic analysis of negative and positive feedback loops for robustness and temperature compensation in circadian rhythms. NPJ Systems Biology and Applications 9 (2023).

40 Avello, P., Davis, S. J., Ronald, J. & Pitchford, J. W. Heat the Clock: Entrainment and Compensation in Arabidopsis Circadian Rhythms. Journal of Circadian Rhythms 17 (2019).

41 Peng, Y.-c., Hasegawa, Y., Noman, N. & Iba, H. Temperature compensation via cooperative stability in protein degradation. Physica A-statistical Mechanics and Its Applications 431, 109–123 (2015).

42 Lim, R., Chae, J. Y., Somers, D. E., Ghim, C.-M. & Kim, P.-J. Cost-effective circadian mechanism: rhythmic degradation of circadian proteins spontaneously emerges without rhythmic post-translational regulation. iScience 24 (2021).

43 Gabriel, C. H., et al. Circadian period is compensated for repressor protein turnover rates in single cells. Proceedings of the National Academy of Sciences of the United States of America 121 (2024).

44 Rosbash, M. The implications of multiple circadian clock origins. PLoS Biol 7, e62, doi:10.1371/journal.pbio.1000062 (2009).

45 Liao, M., et al. The P-loop NTPase RUVBL2 is a conserved clock component across eukaryotes. Nature 642, 165–173, doi:10.1038/s41586-025-08797-3 (2025).

46 Ode, K. L. & Ueda, H. R. Design Principles of Phosphorylation-Dependent Timekeeping in Eukaryotic Circadian Clocks. Cold Spring Harb Perspect Biol 10, doi:10.1101/cshperspect.a028357 (2018).

47 Mosier, A. E. & Hurley, J. M. Circadian Interactomics: How Research Into Protein-Protein Interactions Beyond the Core Clock Has Influenced the Model of Circadian Timekeeping. J Biol Rhythms 36, 315–328, doi:10.1177/07487304211014622 (2021).

48 Nakajima, M., et al. Reconstitution of Circadian Oscillation of Cyanobacterial KaiC Phosphorylation in Vitro. Science 308, 414–415 (2005).

49 Tomita, J., Nakajima, M., Kondo, T. & Iwasaki, H. No Transcription-Translation Feedback in Circadian Rhythm of KaiC Phosphorylation. Science 307, 251–254 (2005).

50 Sugano, S., Andronis, C., Ong, M. S., Green, R. M. & Tobin, E. M. The protein kinase CK2 is involved in regulation of circadian rhythms in Arabidopsis. Proc Natl Acad Sci U S A 96, 12362–12366, doi:10.1073/pnas.96.22.12362 (1999).

51 Uehara, T. N., et al. Casein kinase 1 family regulates PRR5 and TOC1 in the Arabidopsis circadian clock. Proc Natl Acad Sci U S A 116, 11528–11536, doi:10.1073/pnas.1903357116 (2019).

52 Liu, X., et al. FRQ-CK1 interaction determines the period of circadian rhythms in Neurospora. Nat Commun 10, 4352, doi:10.1038/s41467-019-12239-w (2019).

53 Mehra, A., et al. A role for casein kinase 2 in the mechanism underlying circadian temperature compensation. Cell 137, 749–760, doi:10.1016/j.cell.2009.03.019 (2009).

54 Lin, J. M., et al. A role for casein kinase 2alpha in the Drosophila circadian clock. Nature 420, 816–820, doi:10.1038/nature01235 (2002).

55 Meng, Q. J., et al. Setting clock speed in mammals: the CK1 epsilon tau mutation in mice accelerates circadian pacemakers by selectively destabilizing PERIOD proteins. Neuron 58, 78–88, doi:10.1016/j.neuron.2008.01.019 (2008).

56 Tsuchiya, Y., et al. Involvement of the protein kinase CK2 in the regulation of mammalian circadian rhythms. Sci Signal 2, ra26, doi:10.1126/scisignal.2000305 (2009).

57 Beale, A. D., et al. Casein Kinase 1 Underlies Temperature Compensation of Circadian Rhythms in Human Red Blood Cells. J Biol Rhythms 34, 144–153, doi:10.1177/0748730419836370 (2019).

58 Shinohara, Y., et al. Temperature-Sensitive Substrate and Product Binding Underlie Temperature-Compensated Phosphorylation in the Clock. Mol Cell 67, 783–798.e720, doi:10.1016/j.molcel.2017.08.009 (2017).

59 Rothenfluh, A., Abodeely, M. & Young, M. W. Short-period mutations of per affect a double-time-dependent step in the Drosophila circadian clock. Curr Biol 10, 1399–1402, doi:10.1016/s0960-9822(00)00786-7 (2000).

60 Joshi, R., Cai, Y. D., Xia, Y., Chiu, J. C. & Emery, P. PERIOD Phosphoclusters Control Temperature Compensation of the Drosophila Circadian Clock. Front Physiol 13, 888262, doi:10.3389/fphys.2022.888262 (2022).

61 Top, D., Harms, E., Syed, S., Adams, E. L. & Saez, L. GSK-3 and CK2 Kinases Converge on Timeless to Regulate the Master Clock. Cell Rep 16, 357–367, doi:10.1016/j.celrep.2016.06.005 (2016).

62 Cai, Y. D., et al. CK2 Inhibits TIMELESS Nuclear Export and Modulates CLOCK Transcriptional Activity to Regulate Circadian Rhythms. Curr Biol 31, 502–514.e507, doi:10.1016/j.cub.2020.10.061 (2021).

63 Konopka, R. J. & Benzer, S. Clock mutants of Drosophila melanogaster. Proc Natl Acad Sci U S A 68, 2112–2116, doi:10.1073/pnas.68.9.2112 (1971).

64 Lin, J. M., Schroeder, A. & Allada, R. In vivo circadian function of casein kinase 2 phosphorylation sites in Drosophila PERIOD. J Neurosci 25, 11175–11183, doi:10.1523/JNEUROSCI.2159-05.2005 (2005).

65 Grima, B., et al. The F-box protein slimb controls the levels of clock proteins period and timeless. Nature 420, 178–182, doi:10.1038/nature01122 (2002).

66 Edery, I., Zwiebel, L. J., Dembinska, M. E. & Rosbash, M. Temporal phosphorylation of the Drosophila period protein. Proc Natl Acad Sci U S A 91, 2260–2264, doi:10.1073/pnas.91.6.2260 (1994).

67 Zhang, Y., Liu, Y., Bilodeau-Wentworth, D., Hardin, P. E. & Emery, P. Light and temperature control the contribution of specific DN1 neurons to Drosophila circadian behavior. Curr Biol 20, 600–605, doi:10.1016/j.cub.2010.02.044 (2010).

68 Preuss, F., et al. Drosophila doubletime Mutations Which either Shorten or Lengthen the Period of Circadian Rhythms Decrease the Protein Kinase Activity of Casein Kinase I. Molecular and Cellular Biology 24, 886–898 (2004).

69 Nawathean, P. & Rosbash, M. The doubletime and CKII kinases collaborate to potentiate Drosophila PER transcriptional repressor activity. Molecular cell 13 **2**, 213–223 (2004).

70 Nawathean, P., Stoleru, D. & Rosbash, M. A Small Conserved Domain of Drosophila PERIOD Is Important for Circadian Phosphorylation, Nuclear Localization, and Transcriptional Repressor Activity. Molecular and Cellular Biology 27, 5002–5013 (2007).

71 Blau, J. PERspective on PER phosphorylation. Genes & development 22 **13**, 1737–1740 (2008).

72 Price, J., et al. double-time is a novel Drosophila clock gene that regulates PERIOD protein accumulation. Cell 94 **1**, 83–95 (1998).

73 Kloss, B., Rothenfluh, A., Young, M. W. & Saez, L. Phosphorylation of period is influenced by cycling physical associations of double-time, period, and timeless in the Drosophila clock. Neuron 30, 699–706, doi:10.1016/s0896-6273(01)00320-8 (2001).

74 Chiu, J. C., Ko, H. W. & Edery, I. NEMO/NLK phosphorylates PERIOD to initiate a time-delay phosphorylation circuit that sets circadian clock speed. Cell 145, 357–370, doi:10.1016/j.cell.2011.04.002 (2011).

75 Ko, H. W., Jiang, J. & Edery, I. Role for Slimb in the degradation of Drosophila Period protein phosphorylated by Doubletime. Nature 420, 673–678, doi:10.1038/nature01272 (2002).

76 Chiu, J. C., Vanselow, J. T., Kramer, A. & Edery, I. The phospho-occupancy of an atypical SLIMB-binding site on PERIOD that is phosphorylated by DOUBLETIME controls the pace of the clock. Genes Dev 22, 1758–1772, doi:10.1101/gad.1682708 (2008).

77 Konopka, R. J., Pittendrigh, C. & Orr, D. Reciprocal behaviour associated with altered homeostasis and photosensitivity of Drosophila clock mutants. J Neurogenet 6, 1–10, doi:10.3109/01677068909107096 (1989).

78 Szabó, A., et al. The CK2 kinase stabilizes CLOCK and represses its activity in the Drosophila circadian oscillator. PLoS Biol 11, e1001645, doi:10.1371/journal.pbio.1001645 (2013).

79 Edery, I., Zwiebel, L. J., Dembinska, M. E. & Rosbash, M. Temporal phosphorylation of the Drosophila period protein. Proceedings of the National Academy of Sciences of the United States of America 91 **6**, 2260–2264 (1994).

80 Kloss, B., Rothenfluh, A., Young, M. W. & Saez, L. Phosphorylation of period is influenced by cycling physical associations of double-time, period, and timeless in the Drosophila clock. Neuron 30 **3**, 699–706 (2001).

81 Cyran, S. A., et al. The Double-Time Protein Kinase Regulates the Subcellular Localization of the Drosophila Clock Protein Period. The Journal of Neuroscience 25, 5430–5437 (2005).

82 Lee, E. & Kim, E. Y. A Role for Timely Nuclear Translocation of Clock Repressor Proteins in Setting Circadian Clock Speed. Experimental Neurobiology 23, 191–199 (2014).

83 Lin, J.-M., et al. A role for casein kinase 2alpha in the Drosophila circadian clock. Nature 420 6917, 816–820 (2002).

84 Akten, B., et al. A role for CK2 in the Drosophila circadian oscillator. Nature Neuroscience 6, 251–257 (2003).

85 Renn, S. C., Park, J. H., Rosbash, M., Hall, J. C. & Taghert, P. H. A pdf neuropeptide gene mutation and ablation of PDF neurons each cause severe abnormalities of behavioral circadian rhythms in Drosophila. Cell 99, 791–802, doi:10.1016/s0092-8674(00)81676-1 (1999).

86 Stoleru, D., Peng, Y., Nawathean, P. & Rosbash, M. A resetting signal between Drosophila pacemakers synchronizes morning and evening activity. Nature 438, 238–242, doi:10.1038/nature04192 (2005).

87 Shafer, O. T., Rosbash, M. & Truman, J. W. Sequential nuclear accumulation of the clock proteins period and timeless in the pacemaker neurons of Drosophila melanogaster. J Neurosci 22, 5946–5954, doi:10.1523/JNEUROSCI.22-14-05946.2002 (2002).

88 Wang, B., Stevenson, E. L. & Dunlap, J. C. Functional analysis of 110 phosphorylation sites on the circadian clock protein FRQ identifies clusters determining period length and temperature compensation. G3 (Bethesda) 13, doi:10.1093/g3journal/jkac334 (2023).

89 Gardner, G. & Feldman, J. F. Temperature Compensation of Circadian Period Length in Clock Mutants of Neurospora crassa. Plant physiology 68 **6**, 1244–1248 (1981).

90 Singh, S., et al. New Drosophila Circadian Clock Mutants Affecting Temperature Compensation Induced by Targeted Mutagenesis of Timeless. Front Physiol 10, 1442, doi:10.3389/fphys.2019.01442 (2019).

91. Petersen, E. N. et al. The dramatic impact of the PER-DBT interaction on circadian timekeeping and temperature compensation. bioRxiv, 2025.2002.2001.636007, doi:10.1101/2025.02.01.636007 (2025).

92 Smolen, P., Hardin, P. E., Lo, B. S., Baxter, D. A. & Byrne, J. H. Simulation of Drosophila circadian oscillations, mutations, and light responses by a model with VRI, PDP-1, and CLK. Biophysical journal 86 5, 2786–2802 (2004).

93 Narasimamurthy, R., et al. CK1δ/ε protein kinase primes the PER2 circadian phosphoswitch. Proceedings of the National Academy of Sciences of the United States of America 115, 5986–5991 (2018).

94 Masuda, S., et al. Mutation of a PER2 phosphodegron perturbs the circadian phosphoswitch. Proceedings of the National Academy of Sciences of the United States of America 117, 10888–10896 (2019).

