## Supplementary figures and images for "The CK2 and DBT kinases promote temperature compensation of the *Drosophila* circadian clock via distinct pathways"

### Supplemental figures S1 and S2

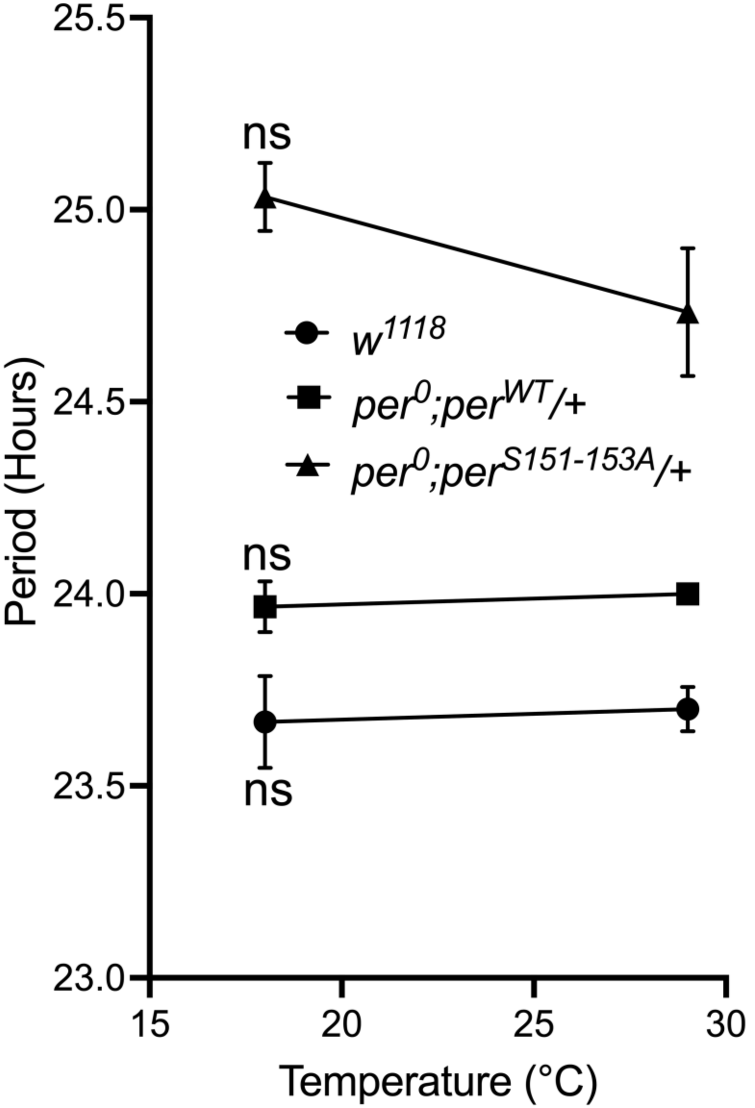

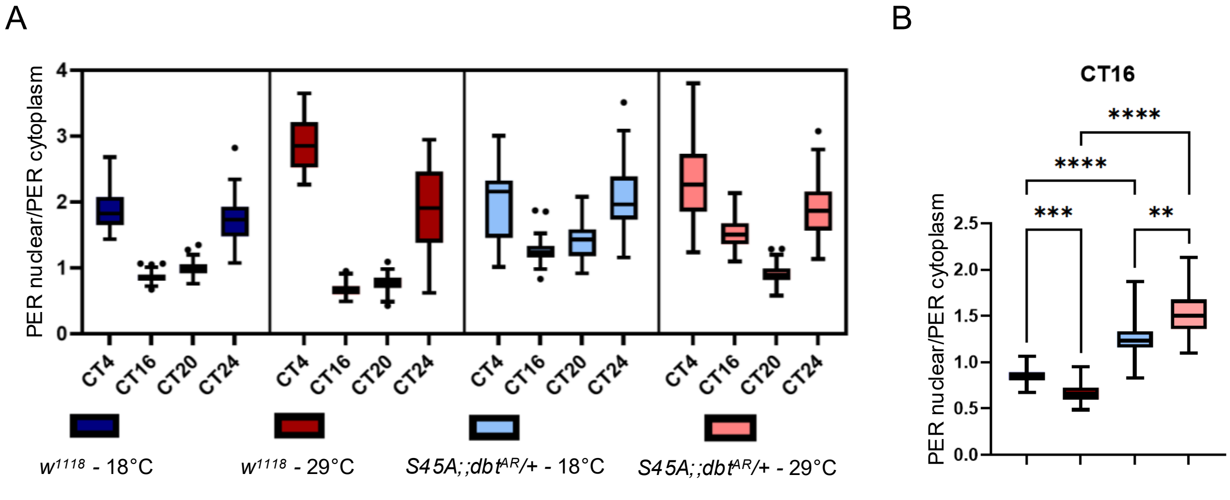
